# Humanized mice bearing CRISPR/Cas9 Disruption of Signal Transducer and Activator of Transcription 1 (STAT1) to Model Primary Immunodeficiency

**DOI:** 10.1101/2022.06.20.496920

**Authors:** Jennifer L. Aron, Timothy Thauland, Humza Khan, Manish J. Butte

**Affiliations:** Department of Pediatrics, Division of Immunology, Allergy, and Rheumatology, University of California, Los Angeles; Los Angeles, CA 90095; USA; Department of Microbiology, Immunology, and Molecular Genetics, University of California, Los Angeles; Los Angeles, CA 90095; USA; California Center for Rare Diseases, Institute for Precision Health; University of California, Los Angeles; Los Angeles, CA 90095; USA

**Author notes:** **Corresponding author:** Manish J. Butte, MD PhD, 10833 LeConte Ave, MDCC Building Room 12-430; Los Angeles CA 90095.

**Keywords:** Primary immunodeficiency, inborn error of immunity, PID, IEI, CRISPR/Cas9, signal transducer and activator of transcription 1, humanized mice, animal model, Janus kinase inhibition

## Abstract

**Background:** The search for a single, pathogenic genetic variant in a patient suspected to have a monogenic inborn error of immunity (IEI) often reveals a multitude of rare variants of unknown significance (VUS). Distinguishing which VUS is disease-causing versus the irrelevant, rare variants from the genetic background is slow and difficult. Advances in gene editing technology, particularly CRISPR/Cas9, promise to accelerate the timeline for the development of single-variant animal models, thus affording an experimental system for validating new genes and their variants.

**Objective:** We sought to demonstrate a proof-of-concept of using CRISPR/Cas9 in human hematopoietic stem cells (hHSC) to develop of humanized mice bearing a hematopoietic deficiency in signal transducer and activator 1 (STAT1).

**Methods:** Using CRISPR/Cas9, we introduced indels into the *STAT1* gene of hHSCs and implanted them into immunodeficient mice. The reconstituted immune systems were assessed by flow cytometry.

**Results:** Mice transplanted with cells edited to eliminate *STAT1* developed human immune systems with diverse cell phenotypes. Lymphocytes from these reconstituted mice showed low expression of STAT1 protein and diminished phosphorylation of STAT1 in response to interferon stimulation. These data mirror the impaired, but not abolished, response to interferons seen in human partial STAT1 deficiency. CRISPR/Cas9 genome editing techniques can be used to rapidly and inexpensively create functional, humanized models of primary immune deficiencies.

## Introduction

Inborn errors of immunity (IEI) include over 500 rare monogenic disorders impacting the development and function of the immune system (Meyts et al. 2020; Tangye et al. 2020, 2021). For patients with suspected IEIs, finding the pathogenic genetic variant is the standard of care today because the cost of sequencing the whole exome (WES) or genome (WGS) has come done and identifying the genetic variant adds significant clinical actionability (Stray-Pedersen et al. 2017). The resulting genomic data are filtered to consider relevant genes and further filtered to consider only the variants that are rare in the population. It is equally likely to find a known pathogenic variant, at which point the diagnostic odyssey comes to an end, as it is to find a “variant of unknown significance” (VUS), at which point the odyssey continues. Approaches to computationally predict the pathogenicity of individual VUS are unreliable at best. Researchers must then choose which VUS is most likely to be causative and embark on the painstaking, costly, and prolonged process of biological validation (Casanova et al. 2014; Rodenburg 2018). Patients with severe symptoms do not always survive the wait for an answer. We propose here a new approach using humanized mice to rapidly and definitively resolve the pathogenicity of unknown variants.

Studying functional defects in primary cells in patients is a tried-and-true approach but incurs the problem that primary cells are often limited and may be difficult to access. For example, in patients with SCID, testing a pathogenic variants may require harvesting bone marrow stem cells. Another problem with working with primary cells is that the “background genome” where common and rare variants can confound interpretation of the putative pathogenic variant of interest. Biochemical rescue experiments on the isogenic background partially resolve this concern, but proving the pathogenicity of takes generating a unrelated cell line to bear the pathogenic variant by reductionistically moving the variant of interest into a new genome. Induced pluripotent stem cells (iPSCs) have emerged as a powerful tool to study a wide array of diseases, including IEIs. Unlike primary cells, iPSCs derived from patient cells offer a nearly infinite source of material on which to experiment, and their pluripotency provides an avenue in which to study the effect of a variant on multiple cell types. Skin cells could be reprogrammed to iPSC and then to hematopoietic stem cells (Pereira et al. 2013), saving a patient from a painful and not-easily-repeatable bone marrow harvest. However versatile, iPSCs in culture can not offer the environmental milieu found in a humanized mouse model that may be needed for testing the role of a gene or variant on IEIs. For example, vaccine responses cannot be tested with reprogrammed iPSCs, but requires a living organism. Moreover, iPSCs derived from the patient do not eliminate the “background genome” problem.

A model that offers biochemical validation would be even more useful if it could help assess potential treatments. Introduction of genetic variants into workhorse cell lines like Jurkat and HEK293 may allow validation of deranged signaling offered by a variant, but do not allow treatments to be tested. Treatment of IEIs with repurposed small-molecule inhibitors or biologicals, and an ideal system would allow simultaneous validation of a variant and testing of potential therapies. The wrong choice of inhibitor can have serious consequences, and there are limited *a priori* ways to know which off-label therapy will work. For example, of two small-molecule inhibitors of PI3K p110δ tested, one inhibitor (leniolisib) halted the disease (Rao et al. 2017), while two others caused significant side effects, including colitis (Coutré et al. 2015; Diaz et al. 2020; Rao et al. 2017). Randomized, head-to-head clinical trials are not feasible when the number of affected patients is very low. An ideal system would allow testing various immunomodulators against various pathogenic variants without exposing patients to potential toxicity.

Gene therapy of IEIs has shown success with a number of diseases (Boztug et al. 2010; De Ravin et al. 2016; Urnov et al. 2005). We envision a day when gene therapy using CRISPR-based gene correction could restore each ultrarare pathogenic variant in IEIs to wildtype. At present, there is no good system for evaluating gene correction on ultrarare children bearing individual, pathogenic variants. Humanized mice offer an opportunity to test variant-specific gene correction before testing on humans. In our mice, we could not only demonstrate the extent to which a single variant contributes to a disease phenotype, but we could similarly check the extent of phenotypic correction after hematopoietic gene editing. For example, one could ascertain whether gene correction in HSCs for NEMO deficiency will correct colitis.

Increased susceptibility to infection is the hallmark of IEI, and diseases caused by variants in the *STAT1* gene are no exception. Partial or complete STAT1 deficiency, or loss-of-function (lof) STAT1, in humans leads to severe bacterial and viral infections. Similarly, patients with *gain*-of-function (gof) STAT1 are typically affected by a trifecta of immunological dysregulation: recurrent infections, autoimmunity, and autoinflammation. Like the majority of IEI, diagnosis of either type of STAT1 variant is hampered by the variability of presenting symptoms (Toubiana et al. 2016), overlapping clinical signs, and lack of sensitive and specific clinical laboratory tests. Without an accurate and timely diagnosis, prompt initiation of optimal treatments remains elusive. Successful hematopoietic stem cell transplants are extraordinarily difficult in STAT1 disease due to inflammation and high rate of secondary graft loss (Leiding et al. 2017). JAK inhibitors are indicated for gof disease, but the choice of inhibitor and the right dosages are still empirically determined for each patient. At this point the diagnosis of lof or gof in STAT1 function can be ascertained from flow cytometric experiments in a research setting (Zimmerman et al. 2019). Because of these reasons, STAT1 offers a good proof-of-concept of the humanized mice approach described here.

## Results and Discussion

To target the *STAT1* gene by CRISPR/Cas9 in human cells, we designed several potential guide RNA sequences for inclusion into ribonucleoproteins (RNPs) and evaluated their cutting efficacy. We employed the T7EI assay or ICE (interference of CRISPR edits) Sanger sequence trace decomposition tool after electroporation. We selected a region including *STAT1* exon 10 in which we observed a variant from a patient in our clinic. One guide sequence (5’-UCCGCAACUAUAGUGAACCU-3’) outperformed other candidates in consistently producing insertions and deletions in human cells, according to T7EI assay or ICE analysis estimates (**Figure 1A-B**). Despite effective electroporation (**Figure 1C**), initial experiments in human HSCs (hHSC) resulted in low levels of insertions and deletions (data not shown). Adding poly(glutamic acid) (PGA) for stabilization of the RNP nanoparticles (Nguyen et al. 2020) and PGE2 for hHSC survival (Genovese et al. 2014; North et al. 2007) enhanced conditions so that levels of indels in hHSCs reached 50% (**Figure 1D**). These data show that under the right conditions, CRISPR/Cas9 can effectively be used to directly target *STAT1* in human HSCs.

**Figure 1.**
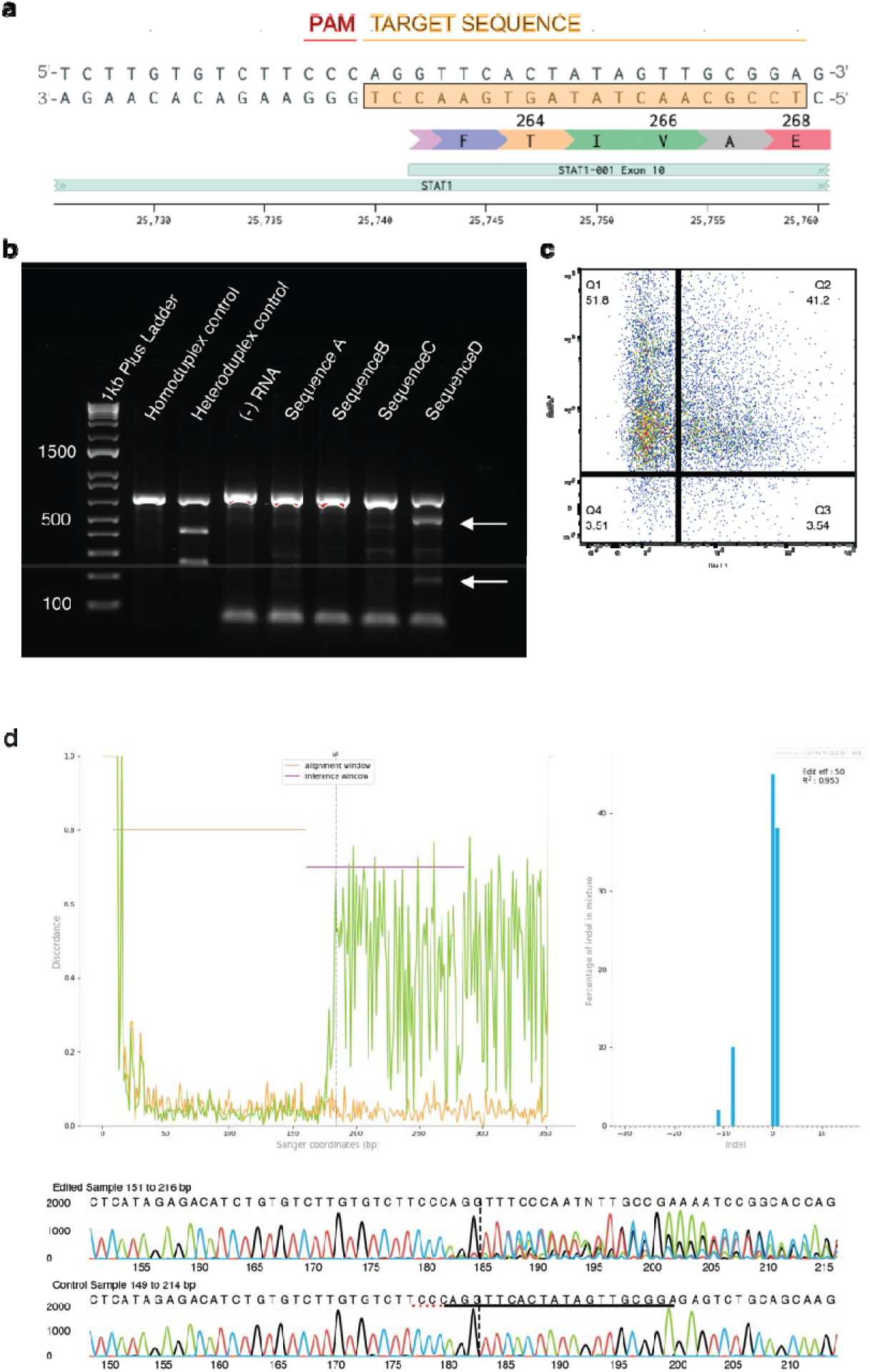
(1.5 columns wide). CRISPR/Cas9-mediated targeting of PID genes in hHSCs. **a**) Gene targeting strategy (sgRNA selection); **b**) H9 cells were electroporated with RNPs containing different guide sequences and control GFP plasmid. T7EI assay on extracted DNA shows fragments that indicate mutated STAT1 by sequence D. **c**) electroporation efficiency; **d**) Indels shown by ICE analysis

Oftentimes, models of human genetic diseases are produced with germline mutations in orthologous murine cells. On the other hand, the immune cells of *humanized* mouse models are human in origin and provide a more useful model. To generate humanized mice, we irradiated immunodeficient neonatal mice (NOD.Cg-*Prkdc*^*scid*^ *Il2rg*^*tm1Wjl*^ *H2-Ab1*^*tm1Doi*^ Tg(HLA-DRB1)31Dmz/Sz) and transplanted them with control or *STAT1*-targeted hHSCs (**Figure 2A**). Both control and experimental mice developed T and B cells and monocytes of human origin in peripheral blood, spleen, and bone marrow (**Figure 2B-C**). Thus, transplantation of control or edited hHSCs into immunodeficient mice results in successful engraftment.

**Figure 2.**
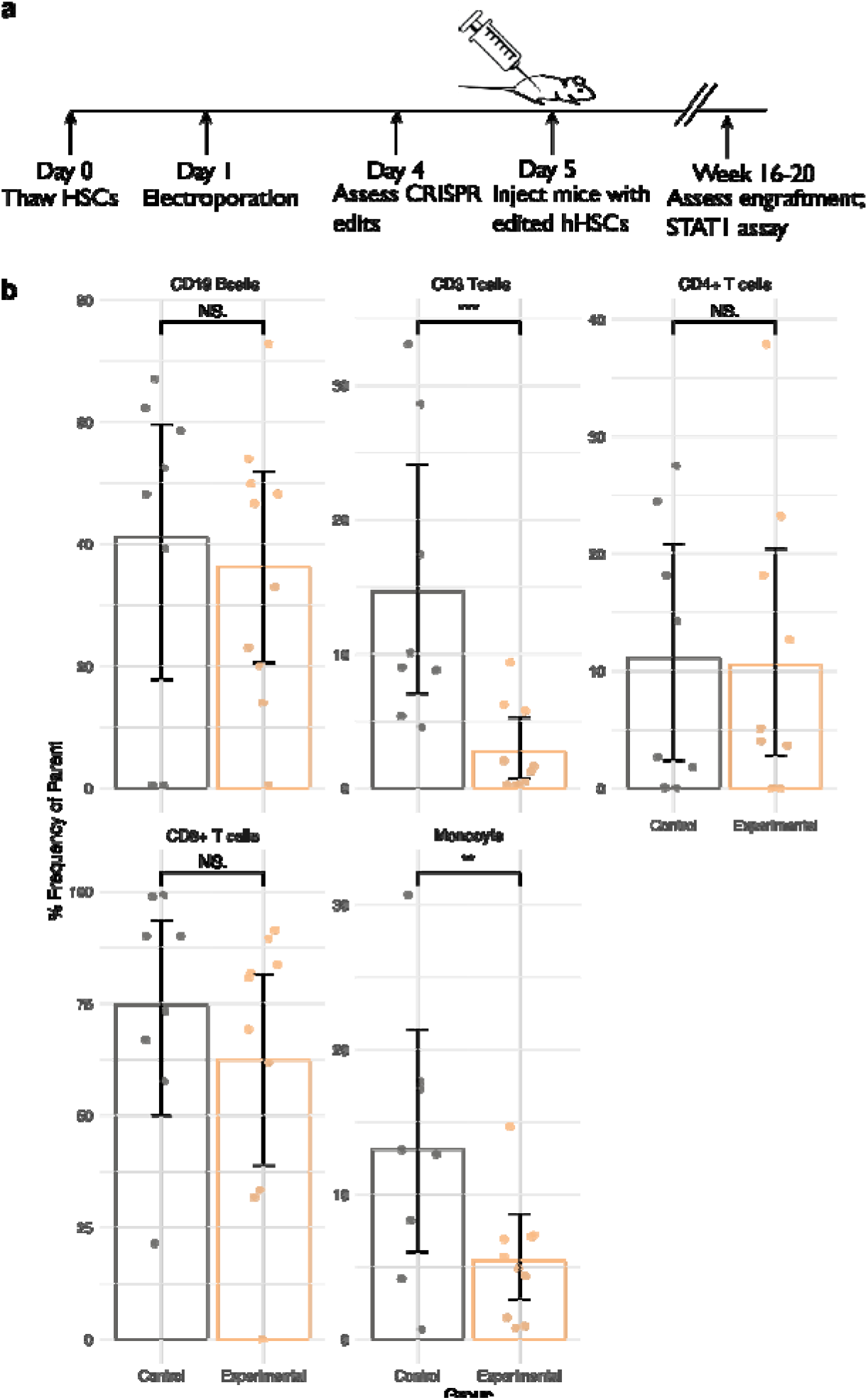
(1.5 columns wide). Generation of a humanized IEI mouse model by implanting edited stem cells into immunodeficient mice. **a)** schematic of the approach; **b)** populations of key immune cells in the transplanted mice.

We next measured the function of STAT1 protein with a phosphorylation assay. Given that interferon receptor signaling activate STAT1 signaling, we evaluated the responsiveness of STAT1 to interferon alpha (IFN-α) and gamma (IFN-γ) in our humanized mice. Peripheral blood mononuclear cells (PBMCs) from mice humanized with edited, STAT1-deficient hHSCs demonstrated diminished total STAT1 (**Figure 3A**). This result confirms that across B cells, T cells, and monocytes, edited humanized mice lack STAT1 expression on the protein level.

**Figure 3.**
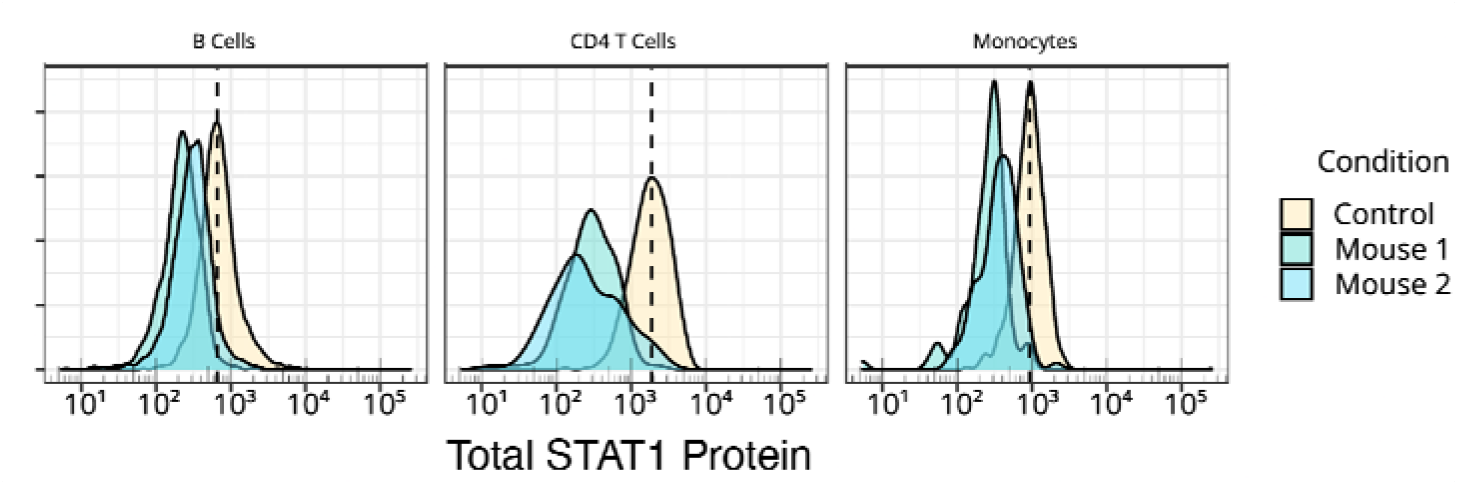
(1.5 columns). CRISPR/Cas9-mediated genetic disruption impairs the expression of STAT1.

Upon stimulation with IFN-α, phosphorylation responses were lower (**Figure 4**). On the other hand, cells derived from unedited hHSCs responded to IFN-α with robust phosphorylation of STAT1. These results show that humanized mice lacking *STAT1* lack appropriate signaling.

**Figure 4.**
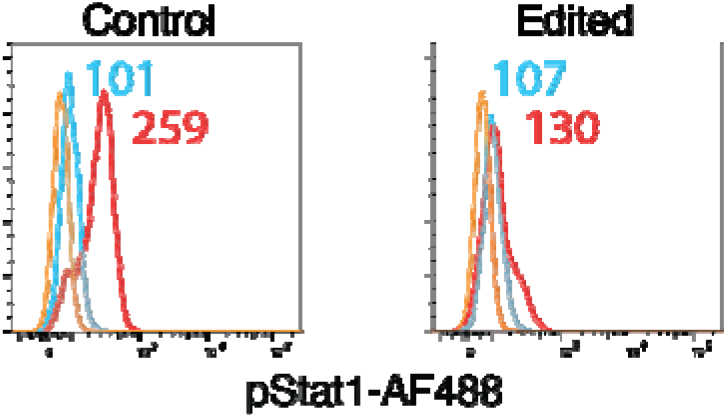
(1 column). CRISPR/Cas9-mediated genetic disruption impairs the function of STAT1 after stimulation with IFN-_α_.

Our humanized mice reconstituted from CRISPR/Cas9-edited HSCs offer a feasible alternative to traditional methods for testing the importance of individual variants of unknown significance. The approach is streamlined so that new rare genetic diseases can be studied in about 20-30 weeks. Humanized murine IEI models can assess the extent to which a single variant contributes to a disease phenotype, can assess the spectrum of infection susceptibility for a new gene, can be used to assess the efficacy of potential repurposed drugs, and can check the extent of phenotypic correction after gene editing. This approach is minimally invasive and alleviates the need for multiple blood draws from patients.

### Clinical implications

There is no standard system for evaluating the pathogenicity of variants of unknown significance in the inborn errors of immunity. Humanized mice reconstituted from Cas9-edited HSCs bearing putative variants provide an opportunity to rapidly test the in vio relevance of genetic variants, test pharmacologic interventions, and trial variant-specific gene correction before testing on humans.

## Acknowledgements

We are grateful to Scott Kitchen’s lab for general advice. We acknowledge funding from the Jeffrey Modell Foundation through a grant from Takeda and NIH/NIAID R01AI153827 to MJB

## Methods

### Mice

NOD-scid IL2rgamma Ab0 Tg (HLA-DR4), NSG-Ab° DR4 compound mutant NSG-Ab° DR4 mice (Jax strain 017637), which lack expression of the murine *Prkdc* gene, the X-linked *Il2rg* gene, and MHC class II while expressing the human leukocyte antigen DR4 gene, were purchased from Jackson Laboratory (Bar Harbor, ME), bred, and maintained under pathogen-free conditions with access to food and water *ad libitum* in the University of California, Los Angeles (UCLA) animal facility. Protocols for animal use were approved by the Institutional Animal Care and Use Committee at UCLA.

### Isolation of human hematopoietic stem cells

Human fetal liver tissue was obtained from Advanced Bioscience Resources (ABR), immediately washed in PBS upon receipt, freed of connective tissue by scalpel, and further homogenized by re-suspension with a 16-gauge blunt needle and 10-mL syringe. The tissue suspension was then incubated with 10 mL of Iscove’s Modified Dulbecco’s Media (IMDM), collagenase (500U/mL), hyaluronidase (2,400 U/mL), DNase (300 U/mL), and penicillin-streptomycin (80 U/mL and 80 mg/mL) for 90 min in a tube rotator at 37 °C, then filtered through a 100 µm cell strainer. Cell density centrifugation media (Ficoll) was laid under the cell suspension, and tubes were centrifuged at 1200 x g for 20 min without brake applied. The buffy coat interface was removed and transferred to tubes with fresh PBS, after which cells were again washed, then subjected to isolation with a human CD34^+^ positive selection kit (StemCell), following the manufacturer’s protocol.

### CRISPR/Cas9 protocol and analyses

Isolated CD34^+^ HSCs were cultured in X-Vivo-10 media (Lonza), IL-3, IL-6, thrombopoietin, stem cell factor [SCF], and Flt3-ligand (PeproTech). RNP was prepared by combining sgRNA (Synthego) with poly(glutamic acid) (PGA) in a 1:0.8 ratio, and complexing with spCas9 (UC Berkeley) at a ratio of 8 µg sgRNA:8µg Cas9 per 100 µL cuvette. In preparation for electroporation, cells were gently spun (at 90 x g), then re-suspended in P3 solution (Lonza), prostaglandin E2 (PGE2), and the RNP-PGA. HSCs were electroporated with the EO-100 program in the 4D-Nucleofector Core Unit after. Loss-of-function cells were produced by electroporation with specific RNP but no donor template, resulting in non-specific insertions and deletions. After a 3-5-day incubation, 5 × 10^5^ HSC cells were ready for transplant. Neonatal immunodeficient NSG Ab0 DR1 mice pups 1-5 days of age underwent conditioning radiation (150 rad of gamma radiation from X-ray irradiator). HSCs were injected into the neonatal liver and pups returned to the dam until weaning.

### HSC engraftment analyses in immunodeficient mice

16 weeks after injection of HSCs, we obtained peripheral blood and confirmed engraftment by flow cytometry to assess reconstitution of T cells and B cells, NK cells, and granulocytes. Analyses were performed in FlowJo.

### STAT1 assay

Peripheral blood mononuclear cells (PBMC) were stained in whole blood for extracellular markers, then washed and stimulated with human IFN-α1 (Cell Signaling Technology, cat. no. 8927SC) or IFN-γ (PeproTech, Inc., cat. no. 300-02) at 10 ng/mL for 20 minutes at 37°C, fixed, permeabilized and stained with anti-human phosphorylated Stat1 Alexa Fluor® 488 mouse anti-Stat1 [pY701], BD Biosciences, cat. no. 612596), total Stat1 (Alexa Fluor® 647 mouse anti-Stat1, BD Biosciences, cat. no. 558560), or isotype controls (BD Biosciences, cat. no. 558055, 558053).

### Statistical Analyses

Data shown are the geometric means plus or minus standard deviations. The statistical analyses were performed using R and RStudio software.

